# The mitochondrial Cu^+^ transporter PiC2 (SLC25A3) is a target of MTF1 and contributes to the development of skeletal muscle *in vitro*

**DOI:** 10.1101/2022.09.05.506690

**Authors:** Cat McCann, Michael Quinteros, Ifeoluwa Adelugba, Marcos N. Morgada, Aida R. Castelblanco, Emily J. Davis, Antonio Lanzirotti, Sarah J. Hainer, Alejandro J. Vila, Juan G. Navea, Teresita Padilla-Benavides

## Abstract

The loading of copper (Cu) into cytochrome c oxidase (COX) in mitochondria is essential for energy production in cells. Extensive studies have been performed with mitochondrial cuproenzymes, such as Sco1, Sco2 and Cox17, which contributes to the metallation of the oxidase. However, limited information is available on the upstream mechanism of Cu transport and delivery to mitochondria, especially through Cu-impermeable membranes, in mammalian cells. The mitochondrial phosphate transporter SLC25A3, also known as PiC2, is also able to bind Cu^+^ and acts as an active copper transporter in eukaryotic cells through these membranes, and ultimately aid in the metallation of COX. We used a well-established differentiation model of primary myoblasts derived from mouse satellite cells, where Cu availability is necessary for growth and maturation, and showed PiC2 is a target of MTF1, its expression is induced during myogenesis and favored by Cu supplementation. PiC2 deletion using CRISPR/Cas9 showed that the transporter is required for proliferation and differentiation of primary myoblasts, as both processes are delayed upon *PiC2* knock-out. The effects of *PiC2* deletion were ameliorated by the addition of Cu to the growth medium, implying the deleterious effects of *PiC2* knockout in myoblasts may be in part due to a failure to deliver sufficient Cu to the mitochondria, which can be compensated by other mitochondrial cuproproteins. Co-localization and co-immunoprecipitation of PiC2 and COX also strongly suggest that PiC2 may act to directly load Cu into COX, which was verified by *in vitro* Cu^+^-transfer experiments. The data indicate an important role for PiC2 in both the delivery of Cu to the mitochondria, COX and, subsequently, the differentiation of primary myoblasts.

## INTRODUCTION

Copper (Cu) is an essential redox cofactor for enzymatic and energy production reactions [1; 2]. Cu can also be toxic at high concentrations and, if deleteriously oxidized, enhances the production of reactive oxygen species [3]. Consequently, cells *must* control Cu levels and charge, which they do via a complex cellular network of transmembrane transporters, soluble chaperones, and transcription factors [4; 5; 6; 7; 8]. These proteins have high binding affinities for Cu, and they simultaneously chelate free Cu in the cells and distribute it to organelles and acceptor proteins [9; 10]. To complete this process, Cu is transferred through direct protein-protein interactions or mechanisms involving ligand exchange [9; 11; 12; 13]. Chelator proteins enable Cu entry into and export out of the cell [13; 14; 15]. The copper transporter 1 (CTR1) facilitates Cu entry to the cell [7; 16]. As Cu enters the cell, it is bound by acceptors, such as glutathione, antioxidant protein 1 (ATOX1), and the Cu-chaperone for superoxide dismutase (CCS) [17]. Some examples of cuproenzymes in eukaryotes are cytochrome *c* oxidase (COX), Cu,Zn-Superoxide dismutase (SOD1), tyrosinase, lysyl oxidase, and peptidyl-glycine-α-monooxygenase. These proteins may be cytosolic, mitochondrial, or secreted via the *trans*-Golgi network. The ATP-driven Cu-transporters ATP7A and ATP7B are responsible for the metallation of the secreted cuproproteins, as they transport cytosolic Cu into the lumen of the trans-Golgi network [5; 6; 9; 18]. Failure in Cu acquisition, distribution, or delivery to appropriate acceptors leads to fatal disorders, such as Menkes’ and Wilson’s diseases. Moreover, cellular respiration depends on the fine-tuning of redox processes, which is finalized by the reduction of O_2_ mediated by COX, where Cu is an essential component for electron transfer.

The skeletal muscle is a contractile tissue composed of multinucleated myofibers formed by the fusion of differentiating muscle cells. Muscle regeneration and development depends on the stem cell population, or satellite cells, located between the muscle sarcolemma and the basal lamina of individual myofibers [19]. Myoblasts proliferation is largely regulated by the transcription factor Pax7 [20; 21; 22; 23; 24; 25; 26; 27; 28; 29; 30], and the chromatin remodeling complex SWI/SNF that promote the expression of *Pax7* in primary myoblasts [25]. Activation of satellite cells requires the expression of the myogenic determination protein (MYOD) and myogenic factor 5 (MYF5) [31; 32; 33; 34; 35]. Myogenic differentiation of these cells is dependent on *Pax7* downregulation, induction of the master regulator *myogenin*, and myoblast fusion to create myofibers [36].

Muscle tissue has an intrinsically high number of mitochondria and, consequently, a high demand for Cu. As such, myogenesis serves as a good physiological model in which to study mitochondrial Cu transport and the role of Cu in mitochondrial biogenesis. Myogenesis results in a number of metabolic and morphological changes that have strong links to Cu biology. For instance, energy production in proliferating myoblasts depends largely on glycolysis, while myotube differentiation requires oxidative phosphorylation, with a subsequent increase in mitochondrial biogenesis and function [37; 38]. Oxidative phosphorylation requires Cu for proper function of COX which as the terminal electron acceptor of the electron transport chain. Myoblasts lacking mitochondrial DNA cannot differentiate [39], and inhibition of mitochondrial protein synthesis or functional impairment causes a failure in myogenesis [40; 41; 42; 43].

It is well-established that Cu is required for the maturation and activity of COX [44; 45]. However, there are still some gaps in our understanding of the pathway responsible for the delivery of Cu across the mitochondrial membranes, and the mechanisms by which it is loaded into COX. In *Saccharomyces cerevisiae*, a non-proteinaceous trafficking system supports mitochondrial Cu transport: a cytosolic Cu ligand (CuL) delivers the ion to the organelle. Therein, the yeast mitochondrial phosphate carrier yPiC2 imports Cu into the matrix, which is eventually utilized by COX and SOD1 [46]. Limited information is available on mitochondrial Cu-transporting systems for other eukaryotes. Sequence homology analyses showed that the mammalian (human and murine) mitochondrial phosphate transporter encoded by *SLC25A3* has a 48% identity to yPiC2. SLC25A3, also called PiC2, is highly conserved in mammals, and it catalyzes the transport of Pi into the mitochondrial matrix [47]. Early homology models suggested that SLC25A3 has six transmembrane (TM) segments, 3-fold symmetry, and N- and C-termini facing the intermembrane space [48]. SLC25A3 has two splice forms: PiC2A, which is expressed exclusively in cardiac and skeletal muscle, and PiC2B, which is ubiquitously expressed [49; 50]. Biochemical analysis of the bovine (bPiC2) transporter showed that the bPiC2A isoform has higher Pi transport affinity than does bPiC2B, whereas bPiC2B has a higher maximal transport rate [51]. Experimental evidence strongly suggests that the principal activity of PiC2 is to supply Pi for oxidative phosphorylation. Mutations in this transporter lead to failure in phosphate transport that impacts muscle function and/or development. No information on additional transporting roles for the mammalian PiC2 homolog have been reported. Despite the importance of Cu to mitochondrial function, the mechanisms by which Cu is transported to the mitochondria are still being elucidated. We hypothesized that in mammalian cells PiC2, plays a similar role to that of its yeast counterpart, yPic2, in transporting Cu^+^ to mitochondria. We used a model of cultured primary myoblasts derived from mouse satellite cells to investigate the role that the murine PiC2 (mPiC2) plays in mitochondrial Cu delivery and skeletal muscle growth and differentiation. We found that mPiC2 is induced during the muscle differentiation process, and in the presence of Cu the expression of the transporter is significantly increased. Bioinformatic analyses of publicly available ChIP-seq datasets [52] for differentiating myoblasts showed that *PiC2* is a target of the metal transcription factor MTF1, and that this binding is increased when the cells are supplemented with Cu [52], which was confirmed by ChIP-qPCR analyses. Homology modeling using Alpha fold showed that PiC2 presents a number of potential metal binding residues in the transmembrane domain. Recombinant PiC2 was expressed and purified to determine the Cu-binding stoichiometry. PiC2A seems to bind two equivalents of Cu, while PiC2B seems to bind three Cu atoms per transporter. In addition, immunohistochemistry and immunoprecipitation analyses showed that mPiC2 interacts with several mitochondrial cuproproteins. *In vitro* Cu transfer experiments showed that PiC2A is able to deliver the ion to COX directly. Finally, CRISPR/Cas9 mediated deletion of *mPiC2* delayed myoblasts growth and differentiation, which may be partially explained by the decreased expression of various mitochondrial cuproproteins in these cells. The proliferation and differentiation defect, as well as deficiency in cuproproteins expression was recovered by Cu supplementation to the culture media, consistent with previous studies [53]. The data strongly suggest that mPiC2 is a novel component of a mitochondrial machinery that supports COX metallation, required for the maturation of the skeletal muscle lineage.

## MATERIALS AND METHODS

### Primary myoblasts culture

Satellite cells were isolated from the leg muscle of 3-6 week-old wild type C57Bl/6 mice as previously described [52]. The tissue was extracted, fragmented, washed with HBSS (Thermo Fisher Scientific, Waltham, MA, USA) and treated with 0.1% Pronase for 1 h at 37°C. The cells were filtered using a 100-μm sieve and resuspended in 3 ml of growth medium containing 1:1 v/v DMEM:F-12, 20% fetal bovine serum (FBS), 5% chicken embryo extract, 25 ng/ml of basic fibroblast growth factor (FGF) and 1% antibiotics. Then, the cells were filtered again using a 40-μm cell sieve and centrifuged at 1000 x *g* for 1 min at room temperature. The cells were separated by Percoll step-gradient (35 and 70%) and centrifuged 20 min at 1850 x *g* at room temperature. The myoblasts located at the lower interface of the 70% Percoll fraction were washed with HBSS, centrifuged 5 min at 1000 x *g*, and resuspended in growth medium for plating.

Samples for proliferating myoblasts were isolated and processed for further experimentation after 48 h of seeing in growth media. To induce differentiation, the myoblasts were seeded on plates coated with 0.02% collagen (Advanced BioMatrix, Carlsbad, CA, USA) [54]. The differentiation treatments are indicated in the figures and were performed as previously described [52; 55]. Briefly, the differentiation medium was supplemented or not with insulin [52; 55; 56; 57]. Supplementation with CuSO4 or the chelator tetraethylenepentamine (TEPA) was done at 100 μM for proliferating and 30 μM for differentiating myoblasts as previously established [52; 55].

HEK293T cells were purchased from ATCC (Manassas, VA) and were maintained in growth media containing Dulbecco’s modified Eagle’s medium (DMEM) supplemented with 10% fetal bovine serum (FBS) and 1% penicillin-streptomycin in a humidified incubator at 37 °C with 5% CO_2_.

### Plasmid construction, virus production, and transduction of primary myoblasts

CRISPR/Cas9 plasmid construction was performed by three custom-designed sgRNAs to recognize the intron/exon junction 2 of the *Slc25a3 (Pic2*) murine gene (Reference Sequence: ENSMUSG00000061904). Each gRNA consisted of 20 nucleotides complementary to the sequence that precedes a 5’-NGG protospacer-adjacent motif (PAM) located in the targeted intron. Specificity was validated by a search through the entire genome to avoid off-target effects. The sequence of gRNAS designed were: PiC2-gRNA2_Intr_F 5’-CACCGCAACAATACAAACCTGCATG-3’ and PiC2-gRNA2_Intr_R 5’-AAACCATGCAGGTTTGTATTGTTGC-3’. Preparation of CRISPR/Cas9 lentiviral constructs was performed using the lentiCRISPRv2 oligo cloning protocol [58]. Briefly, sense and antisense oligos obtained from Integrated DNA Technology (IDT), were set according to the designed sgRNA and were annealed and phosphorylated to form double stranded oligos. Subsequently, they were cloned into the BsmBI–BsmBI sites downstream from the human U6 promoter of the lentiCRISPRv2 plasmid [58; 59] (kind gift from Dr. F. Zhang; Addgene plasmid # 52961). The empty plasmid that expresses only Cas9 but no sgRNA was included as a null knockout control.

To generate Lentiviral particles, 5 x 10^6^ HEK293T cells were plated in 10 cm dishes. The next day, transfection was performed using 15 μg of the sgRNA-containing CRISPR/Cas9 constructs mixed with the packing vectors pLP1 (15 μg), pLP2 (6 μg), pSVGV (3 μg). Transfections were performed using Lipofectamine 2000 according to the manufacturer’s instructions (Invitrogen). The media was changed the next day to 10 ml DMEM (Life Technologies) supplemented with 10 % FBS (Life Technologies). The viral supernatant was harvested after 24 and 48 h of incubation and filtered through a 0.22 μm syringe filter (Millipore). To infect primary myoblasts, 5 ml of the filtered supernatant supplemented with 8 μg/ml polybrene (Sigma) were used to transduce two million cells. After overnight incubation, infected cells were then selected in growth media containing 2 μg/ml puromycin (Invitrogen). Stable myoblasts were maintained in growth media containing 1 μg/ml puromycin.

Parallel 1 plasmids encoding the human *Pic2* isoforms A and B labeled with an N-terminus hexa-histidine tag for protein expression were a generous gift of Dr. Paul Cobine (Department of Biological Sciences, Auburn University) and were previously reported [53; 60].

### Antibodies

The primary antibodies used were rabbit anti-GAPDH (A19056), anti-PiC2 (custom made against the 38-47 residues “GQPRRPRNLA” from the human protein, which cross reacts with the murine protein), anti-SCO1 (A6734), anti-Sco2 (A7051) and the mouse anti-His-Tag (AE003) were from Abclonal Technologies (Woburn, MA). The mouse anti-MTF-1 (H-6, sc-365090), anti-sarco/endoplasmic reticulum Ca^2+^-ATPase (SERCA, sc-271669), anti-COX2 Antibody (D-5, sc-514489) were from Santa Cruz Biotechnologies (Dallas, TX). The mouse anti-complex I antibody (18G12BC2) was from Thermo Fischer Scientific. Hybridoma supernatants against Pax7, anti-myosin heavy chain (MF20) and Myogenin (F5D) were obtained from the Developmental Studies Hybridoma Bank (University of Iowa; deposited by A. Kawakami, D. A. Fischman and W. E. Wright, respectively).

The secondary HRP-conjugated anti-mouse and anti-rabbit antibodies were from Invitrogen (31430 and 31460, respectively) and the fluorescent goat anti-rabbit Alexa-488 and anti-mouse Alexa-594 secondary antibodies were from Thermo Fisher (A-11008 and A-21203, respectively).

### Primary myoblast immunofluorescence

Primary myoblasts for immunofluorescence were grown on glass bottom Cellview Advanced TC culture dishes (Greiner Bio One). Samples were obtained for proliferation and at 24, 48, and 72 h after induction of differentiation. Cells were fixed in 10% formalin, permeabilized with PBT buffer (0.5% Triton-X100 in phosphate buffered saline [PBS]) and blocked in 5% horse serum in PBT. Cells were incubated with the indicated primary antibodies (1:100) in blocking solution overnight at 4°C. The samples were then washed three times with PBT solution for 10 min at room temperature. Then, the cells were incubated with the goat anti-rabbit Alexa-488 secondary antibody (1:500) in blocking solution for 2 h at room temperature and 30 min with DAPI. Cells were counterstained with DAPI and imaged with a Leica TCS SP5 Confocal Laser Scanning Microscope (Leica) using a 40X water immersion objective.

### Gene expression analyses

Three independent biological replicates of proliferating and differentiating primary myoblasts were washed with ice cold PBS and then the RNA extracted using Trizol (Invitrogen). cDNA synthesis was performed using 1 μg of RNA, DNase I amplification grade (Invitrogen 18068-015), and Superscript III (Invitrogen 18080-400) according to the manufacturer’s instructions. Changes in gene expression were analyzed by quantitative RT-PCR using Fast SYBR-Green master mix (Thermo Fisher Scientific) on the ABI StepOne Plus Sequence Detection System (Applied Biosystems) using the comparative Ct method [61] and *Ef1α* as a control. The primers used for *mPic2:* forward 5’-CTGGTGCACGATGGCCTG-3’ and reverse 5’-AACCAAGCTGTATGTGT-3’; *Ef1α:* forward 5’-AGCTTCTCTGACTACCCTCCACTT-3’ and reverse 5’-GACCGTTCTTCCACCACTGATT-3’.

### Western blot analyses

Proliferating and differentiating primary myoblasts were washed with PBS and solubilized with RIPA buffer (10 mM PIPES, pH 7.4,150 mM NaCl, 2 mM EDTA, 1% Triton X-100, 0.5% sodium deoxycholate, and 10% glycerol) containing Complete Protease Inhibitor. Lysates were then sonicated ten times on high power for 30 sec and left to rest for 30 sec. Protein concentrations were determined using Bradford assay [62]. Twenty micrograms of protein were separated on 10% SDS gels and then transferred to PVDF (Millipore) membranes. Membranes were blocked for 1 h at room temperature in blocking buffer (Biorad) and then incubated overnight in the indicated primary antibodies diluted 1:1000 in PBS. Membranes were washed three times in PBST and then incubated for 2 h at room temperature in the indicated secondary antibodies diluted 1:1000 in PBS. Following two washes in PBST and one wash in PBS, membranes were treated with horseradish peroxidase (HRP) substrate for enhanced chemiluminescence (ECL; Tanon), and then imaged on an Analytik Jena imager. GAPDH was used as a loading control.

### Chromatin immunoprecipitation analyses

Three independent biological replicates of proliferating and differentiating primary myoblasts were cross-linked with 1% formaldehyde and incubated for 10 min at room temperature with continuous mild shaking. Glycine was used to inactivate the fixative reagent and cells were incubated for 5 min at room temperature. Samples were washed 3 times with 10 ml of ice-cold PBS supplemented with Complete Protease Inhibitor (Roche, Basel, Switzerland) and cells were resuspended in 1 ml of ice-cold PBS supplemented with Complete Protease Inhibitor. Samples were centrifuged for 5 min at 5000 g at 4°C and the PBS discarded. Primary myoblasts were lysed using the SimpleChIP Plus Sonication Chromatin IP Kit (Cell Signaling Technology), following the manufacturer’s instructions. Samples were sonicated as previously described [52]; proliferating cells were sonicated 3 times for 5 min, 30 s by 30 s at mild intensity and 5 times for differentiating myoblasts using a Bioruptor UCD-200 (Diagenode, Denville, NJ, USA). Then the samples were incubated with the anti-MTF1 antibody or IgG as a negative control[52]. Immunoprecipitated material was collected with magnetic beads, and washed with 1x ChIP buffer supplemented with NaCl, as recommended by the manufacturer. Samples were eluted in 1x elution buffer, incubated overnight at 65 °C and reverse crosslinked with NaCl. The resulting DNA was purified using the ChIP DNA clean concentrator, following the manufacturer’s instructions (Zymo Research, Irvine, CA, USA). The DNA was stored at −80°C until further analysis by semiquantitative real-time PCR (qPCR). The primer sequences in the promoter region of *mPiC2* were: forward 5’-GGACGTCATCGCGTCCTC-3’ and reverse: 5’-AGGTGACCTACTCACTCT-3’.

The ChIP-seq data analyzed here for PiC2 was previously made publicly available with the GEO accession number: GSE116331, and can be downloaded here: https://www.ncbi.nlm.nih.gov/geo/query/acc.cgi?acc=GSE116331

### Sequence analyses and modeling of the putative metal binding sites of PiC2 (SLC25A3)

Protein sequence analyses were performed using the murine and human sequences for PiC2. These were aligned with MUSCLE [63] and ESPript software ([64; 65]). Due to the absence of a crystal structure for the transporter, we used Alpha Fold structure prediction software [66; 67] to obtain a model of the human PiC2 protein (Uniprot accession number: A0A024RBE8). The conserved putative copper binding residues in this model are highlighted in green.

### Cloning, expression, and purification of proteins

Human Pic2A and Pic2B were previously cloned [53] into the pHis-Parallel1 vector which adds a N-terminal 6His-tag [60] and then transformed into *Escherichia coli* BL21 (DE3) competent cells. Protein expression was performed followed the autoinducing media protocol [68]. Purification of Pic2 proteins followed an established membrane protein purification procedure using a Ni-NTA column to elute the His-tagged proteins [14; 69; 70; 71; 72; 73]. Briefly, purification steps were carried out between 0 - 4 °C. Cells were suspended in buffer A (25 mM Tris, pH 7.0, 100 mM sucrose, 1 mM phenylmethylsulfonyl fluoride (PMSF; Sigma) and disrupted with a French press at 20,000 p.s.i. Cell debris was removed by centrifugation at 8,000 x *g* for 20 min at 4 °C. To pellet the membranes, the supernatant was then centrifuged at 229,000 x *g* for 1 h at 4 °C, and resuspended in buffer A at a concentration of 10-15 mg/ml. Membrane proteins were diluted and solubilized to a final concentration of 3 mg/ml in buffer B (25 mM Tris, pH 8.0, 100 mM sucrose, 500 mM NaCl, 1 mM PMSF) containing 0.75% dodecyl-β-D-maltoside (DDM; Calbiochem). Then incubated for 1 h at 4 °C with mild agitation and centrifuged at 229,000 x *g* for 1 h. The supernatant was incubated at 4 °C overnight with Ni^2+^-nitrilotriacetic acid resin (Qiagen) pre-equilibrated with buffer B, 0.05% DDM, and 5 mM imidazole. Next day, the samples were washed with buffer B, containing 0.05% DDM and 20 mM imidazole, and the protein was eluted with buffer B, 0.05% DDM, 250 mM imidazole. Fractions were concentrated and buffer exchange was performed using 10 kDa cut-off centricons (Millipore) to final concentrations of 25 mM Tris, pH 8.0, 100 mM sucrose, 50 mM NaCl, and 0.01% DDM (buffer C).

A stable chimera encoding for cytochrome c oxidase (apoCuA) was expressed and purified by chromatography as previously described [45; 74]. Briefly, a 10 ml culture of BL21(DE3)-pET-9a/CuATt3L was grown overnight in LB medium, supplemented with 50 μg/ml of kanamycin at 37°C. This bacterial culture was used to inoculate 5-6 L cultures and allowed to grow at 37°C for 2-2.5 h until reaching an OD600 = 0.8. Protein expression was induced by adding 1 mM Isopropyl b-D -thiogalactopyranoside (IPTG) and incubated at 37 °C for 5-6 h or 30°C overnight. Bacteria was then pelleted 5000 rpm for 15 min and resuspended in 30 ml of Tris-HCl 50 mM, pH 8.0 supplemented with 2 mM PMSF, 40 μl of DNAse (stock 10 mg/ml), and 5 mM MgCl2. Then, cells were lysed by sonication (5 pulses of 30 seconds each, with 1 min pauses) at the maximum power should be enough. Samples were heat precipitated at 55-60 °C for 10-15 min until precipitate appeared, and centrifuged at 20,000 rpm, at 4°C. Supernatants were collected and DNA was further precipitated by adding cold streptomycin sulfate. Samples were centrifuged again at 20000 rpm, at 4°C and supernatant collected for further fractionation with ammonium sulfate as described. After another centrifugation step at 13,000 rpm for 5 min at 4°C the pellet containing the protein was resuspended in 5 ml of Tris 50 mM pH 8.0. A final precipitation of remaining cell debris was performed by adding 1 ml of ice cold 5 M sodium acetate pH 8.0 every 10 ml of extract and centrifugation at 13000 rpm for 10 min at 4°C. A neutralization step follows by using 100 μl of 5 M NaOH for every 1ml of sodium acetate added. Samples were then dialyzed to eliminate salts by adding 5mM DTT and 5 mM EDTA against Kpi 50 mM pH 7.5. The next day, the proteins are collected from the dialysis bag and further purified by chromatography on a Q-sepharose Fast Flow column in 50 mM Kpi pH 7.5 containing 1 mM DTT, by collecting the CuA protein in the flowthrough. Proteins were aliquoted and stored in buffer containing 100 mM Pi pH 6.0, 100 mM KCl, 1mM DTT at −20°C until further use. All protein concentrations were determined using Bradford assay [62].

### Cu^+^ loading to PiC2 and metal binding analyses

Cu^+^ loading was performed by incubating 20 μM of each apo-protein in the presence of a 10 M excess of CuSO4, 25 mM Hepes (pH 8.0), 150 mM NaCl, and 10 mM ascorbate for 10 min at RT with gentle agitation, as previously described [14; 75]. The unbound Cu^+^ was removed by washing the samples in 10 kDa cut-off centricons. Protein concentrations were determined using Bradford assay prior to mineralization of the samples using 35% HNO_3_ (trace metal grade) for 1 h at 80°C followed by neutralization using 3% H_2_O_2_. Cu bound to the protein was measured by atomic absorption spectrometry (AAS). Stoichiometry of Cu^+^ bound to the PiC2A and Pic2B proteins was calculated by normalizing Cu^+^ content to protein concentration and then normalizing holo-protein Cu/protein ratios and subtracting the background of the apo-protein Cu/protein ratios.

### Ultra-trace copper analysis method

The method for copper ultra-trace (< 1 ppm) analysis of all samples was adapted from previously described protocols [76; 77; 78; 79; 80]. Briefly, Cu quantification was carried out using an atomic absorption spectrometer (AAS) PerkinElmer AAnalyst 800 with a Cu hollow cathode lamp as the radiation source. The AAS was equipped with a graphite furnace (GF-AAS) and UltraClean THGA^®^ graphite tubes (PerkinElmer). This technique allows for accurate ultra-trace copper analysis with limited volume samples, minimizing dilution of samples. For accurate and contaminant-free measurements, all analytical glassware and consumables were acid washed overnight in 5% (v/v) hydrochloric acid and rinsed with 18 MΩ purified water before use [76; 77; 78; 79; 80]. Cu standard solutions were prepared from a 1000 mg/L solution (Sigma-Aldrich) to obtain a calibration curve and determine the dynamic range of the method. The limit of detection for Cu, computed as three times the standard deviation of the intercept of the calibration (3σ), was 10 ppb, and the limit of linearity (LOL) was established at 300 ppb. In a typical analysis of a sample, a known mass of sample was digested in concentrated nitric acid using single-stage digestion [76; 77; 78; 79; 80]. The resulting solution was analyzed for copper in the AAS with measurements carried out at least in triplicates. Cu content on each sample was normalized to the initial mass of protein per sample.

### Cu^+^ transfer experiments

Cu^+^ transfer from the His-tagged metallated holo-PiC2A to apoCuA was performed similarly to previous experiments using Cu^+^-ATPases, with minor modifications [14; 15; 72; 75]. Briefly, PiC2A was loaded or not with Cu^+^, as described above, and bound to a Ni^2+^-nitrilotriacetic acid column in buffer. Apo-CuA was used as Cu^+^ acceptor. Proteins were allowed to interact for 15 min at room temperature and proteins were separated by washing with 25 mM Hepes pH 8.0, 100 mM sucrose, 500 mM NaCl, 0.01 % DDM, 0.01 % azolectin, 10 mM ascorbic acid, 20 mM imidazole followed by elution with 25 mM Hepes pH 8.0, 100 mM sucrose, 500 mM NaCl, 0.01 % DDM, 0.01 % azolectin, 10 mM ascorbic acid, and 300 mM imidazole. Cu^+^ content was determined by the Ultra-trace copper analysis method as described above. Controls were performed where apo-PiC2 was incubated with apoCuA and were subjected to the same procedure.

### Immunoprecipitation

Proliferating and differentiating primary myoblasts grown in the presence or absence of CuSO4 were washed 3 times with ice-cold PBS and resuspended in freshly made IP lysis buffer (50 mM Tris-HCl, pH7.5, 150 mM NaCl, 1% Nonidet P-40, 0.5% sodium deoxycholate, and complete pro-tease inhibitor). Cell extracts were incubated with the anti-PiC2 primary antibody at 4 °C for 2 h, followed by an overnight incubation with Pure-Proteome Protein A/G mix magnetic beads (Millipore Sigma). Samples were washed as indicated by the manufacturer, and immunoprecipitated proteins were eluted in freshly prepared IP-elution buffer (10% glycerol, 50 mM Tris-HCl, pH 6.8, and 1 M NaCl) at room temperature for 1 h, as previously described [81]. Samples were analyzed by western blot probing for the antibodies indicated in the figures.

### Metalloproteomic analyses

Analysis of cuproproteins of differentiating murine myoblasts was performed by 2-dimension grazing-exit x-ray fluorescence (2D-GE/XRF) coupled to liquid chromatography-mass spectrometry (LC-MS/MS) as previously described [82; 83]. Briefly, 100 μg of whole cell extract of primary myoblasts differentiated for 24 h in the presence or absence of insulin and Cu were resolved in 10% native polyacrylamide gels. Denaturing agents were not used for sample preparation or electrophoresis. Gels were blotted onto PVDF membranes using a wet transfer system and blots were analyzed using synchrotron micro-XRF at the GSECARS beamline 13-ID-E, at the Advanced Photon Source (APS), Argonne National Lab (Illinois, USA). The incident X-ray beam energy was tuned to 10.2 keV for these analyses, and focused to a relatively broad spot size of ~ 150×200 μm using rhodium-coated silicon mirrors in a Kirkpatrick-Baez geometry [84]. Energy dispersive X-ray fluorescence spectra was collected using a four element silicon drift detector (Vortex ME4, SII NanoTechnology). Spectra were collected in mapping mode (20 mm x 70 mm maps) by raster scanning the beam through the incident beam in a continuous scan mode so that maps have a 250 μm pixel size with an accumulation time of 80 ms per pixel. Maps of total measured Cu Kα fluorescence intensity were generated, normalized to incident flux, using the GSECARS Larch software [85]. PVDF membranes were marked etched into the edges of the blot and the images obtained using micro-XRF mapping were superimposed to the membranes. The bands corresponding to the Cu signal were excised for tryptic digestion and identification by LC-MS/MS at the University of Massachusetts Chan Medical School, Proteomics and Mass Spectrometry Facility. Data was analyzed with the software Scaffold_3.5.1 (Proteome Software Inc.).

## RESULTS

### Expression of *mPiC2* is induced during primary myoblast differentiation

Differentiating myoblasts have a high demand for Cu and, as such, require an increase in the levels of imported Cu and mobilized within the cell [52; 55]. As a means of assessing whether mPiC2 may play a role in Cu transport during myogenesis, its expression and distribution in differentiating primary myoblasts was assessed (**Fig. 1**). Confocal microscopy analysis revealed a punctate cytosolic staining pattern for mPiC2, that was preserved from proliferating myoblasts to differentiated myotubes (**Fig. 1A**). Western blot analyses show mPiC2 expression was also upregulated in response to the induction of differentiation. *mPiC2* mRNA levels increased 25-fold in day 1 differentiated myoblasts compared to proliferating controls (**Fig. 1B**). This increased further to approximately 150-fold higher than controls at days 2 and 3. A similar increase in mPiC2 protein was also observed, with the highest protein levels present at 72 h after inducing differentiation **(Fig. 1C**). These data show that PiC2 expression is induced during differentiation, suggesting that it may be a relevant player during myogenesis.

**Fig. 1.**
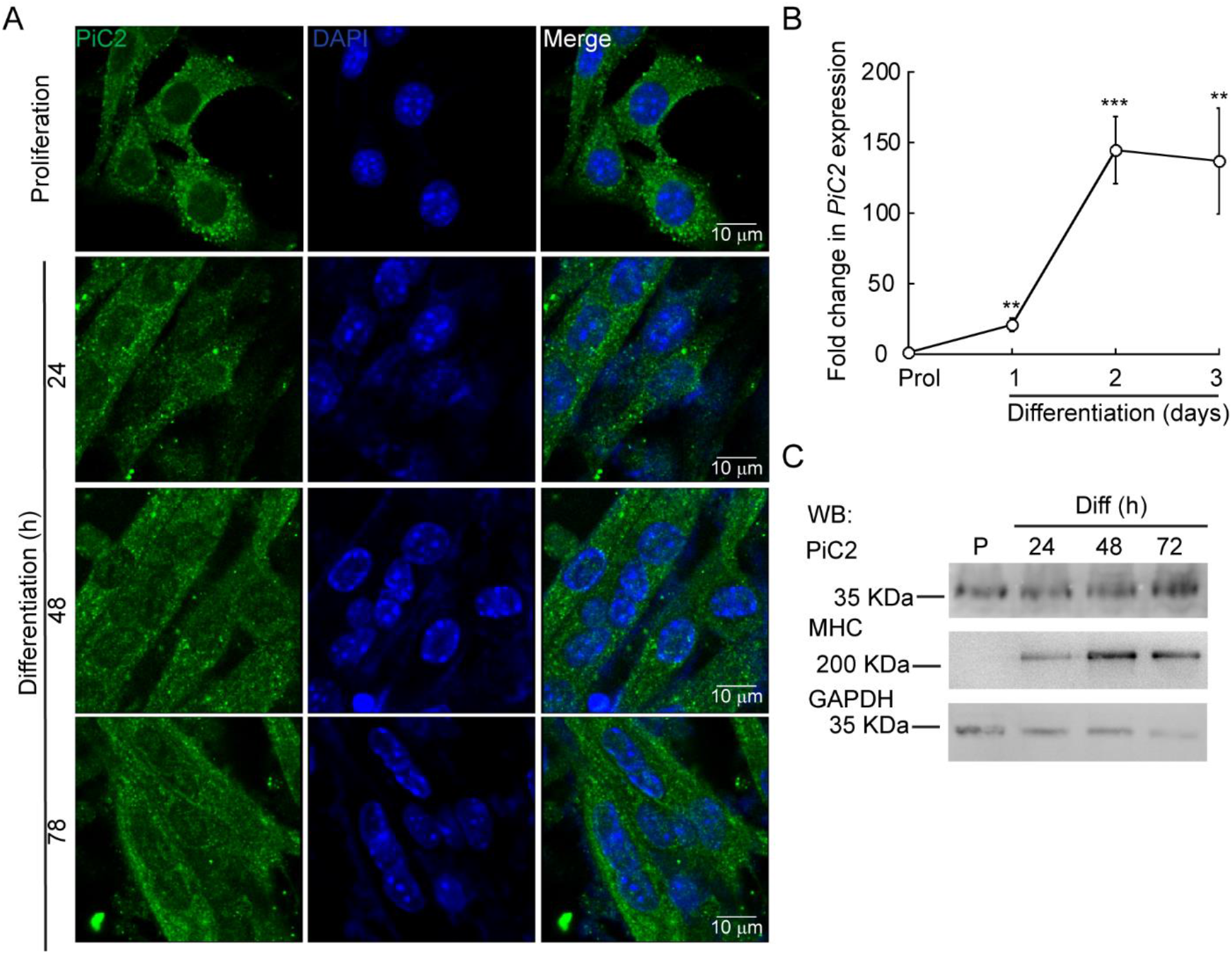
Expression of mPiC2 increases over the course of differentiation in primary myoblasts derived from mouse satellite cells. **(A)** Confocal microscopy analysis of PiC2 expression and localization. Proliferating (48 h) and differentiating (24, 48, and 72 h) primary myoblasts were fixed, and stained with a fluorescent antibody against PiC2 (green) and DAPI (blue). **(B)** *mPiC2* mRNA expression is induced during differentiation. qRT-PCR analysis of *mPiC2* mRNA levels. Values normalized to *Ef1α* and non-differentiated (Prol) controls, *n*=3. **(C)** mPiC2 protein expression is induced during differentiation. Representative western blot of PiC2 protein levels. Membranes were blotted with PiC2, the differentiation marker myosin heavy chain (MHC), and GAPDH was used as loading control. All values are reported as means ± SE. Significance was determined by two-way ANOVA; ** *p*<0.01 and *** *p*<0.001 compared to Prol cells.

### *mPiC2* expression is controlled by MTF1 binding and it’s enhanced by Cu in differentiating primary myoblasts

Cultured primary myoblasts can be differentiated via serum deprivation and insulin supplementation. In the absence of insulin, the myoblasts differentiate poorly [86]. However, differentiation can be restored by the addition of Cu to the growth medium. By contrast, Cu chelation by TEPA impaired differentiation even in cells grown in the presence of insulin [55]. Moreover, Cu has been shown to induce the expression of lineage specific genes, such as *Pax7, myogenin, myosin heavy chain*, and thereby promotes the growth and differentiation of cultured primary myoblasts [55]. To determine whether *mPiC2* expression was impacted by cellular Cu levels, primary myoblasts were treated with TEPA and/or non-toxic concentrations of Cu in the presence or absence of insulin in the culture medium (**Fig. 2**). Myoblasts grown in the presence of insulin and the absence of Cu (black) are considered the normal differentiation control. *PiC2* expression increases upon differentiation, with mRNA levels at day 3 roughly 150-fold higher than those observed in proliferating (Prol) cells (**Fig. 2A**). Cells grown in media depleted of insulin (gray), by contrast, show a modest rise in *mPiC2* expression level compared to Prol control cells. Notably, Cu treatment in the absence of insulin (green) is sufficient to induce *mPiC2* expression to a level even higher than that of the normal differentiation control: roughly 300-fold higher than the Prol cells at day 3 of differentiation compared to 150-fold higher, respectively. Conversely, treatment with TEPA (pink) decreased *mPiC2* expression compared to the differentiation control, with the levels at day 3 of differentiation on par with that of cells grown in the absence of both insulin and Cu. Cu treatment following TEPA treatment (blue) rescues the phenotype, with *mPiC2* expression levels approximately the same as those observed in the differentiation control.

**Fig. 2.**
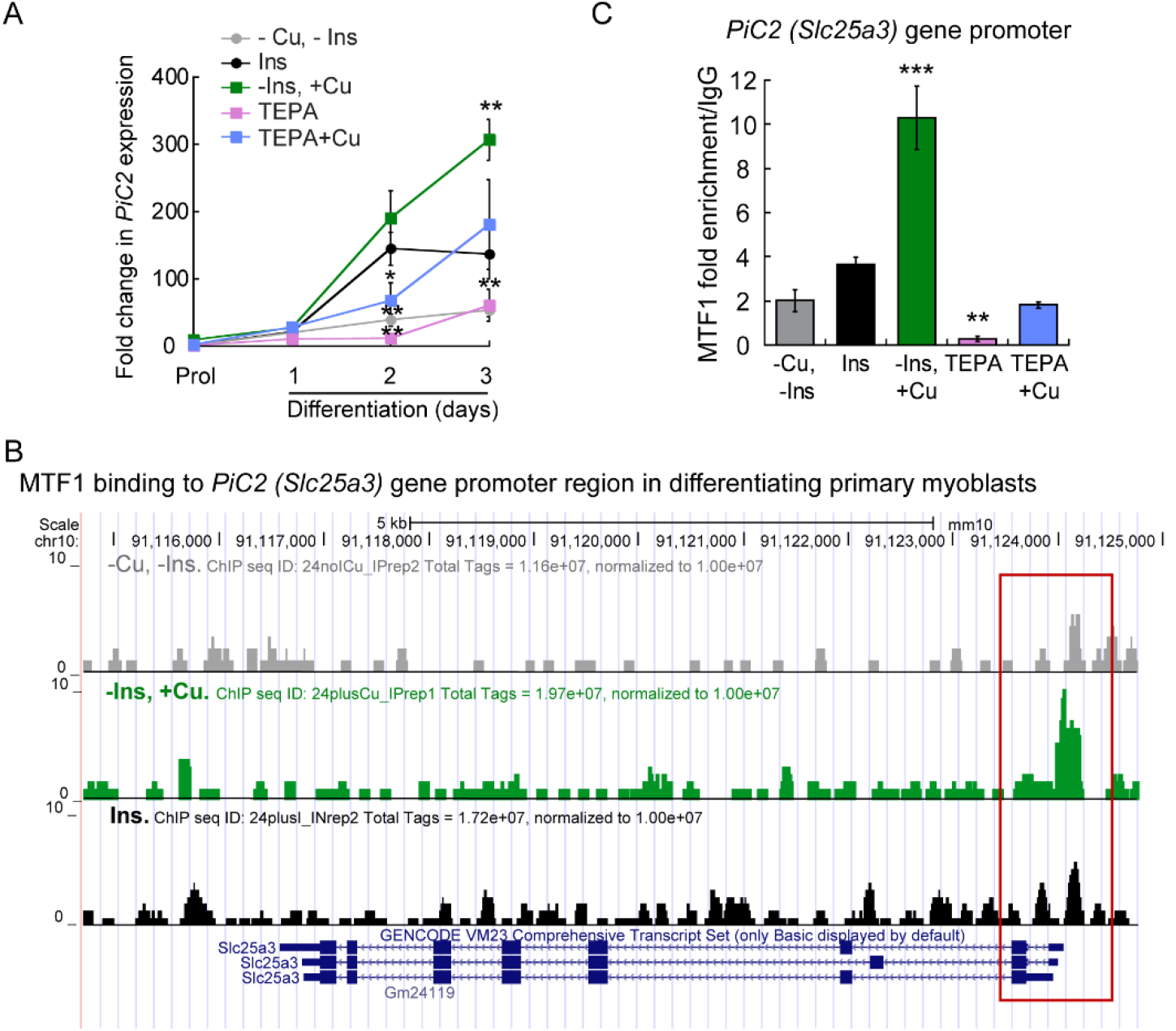
*mPic2* is a target gene of MTF1 in differentiating primary myoblasts, and Cu promotes this binding. **(A)** mRNA levels of *mPiC2* in proliferating and differentiating primary myoblasts. *mPic2* expression was measured by qPCR. Values normalized to *Ef1α* and non-differentiated (Prol) controls, *n=3*. **(B)** Genome browser tracks of ChIP-seq experiments assessing MTF1 binding to the *mPic2* promoter. Promoter region of *mPic2* shown in red box. **(C)** ChIP-qPCR validation of MTF1 binding the *mPic2* promoter region. MTF1 enrichment at the *mPic2* promoter was assessed by ChIP-qPCR, *n*=3. All values are reported as means ± SE. Significance was determined by two-way ANOVA; ** *p*<0.01 and *** *p*<0.001 compared to cells differentiated with insulin.

In conjunction with Cu, the metal-sensing transcription factor MTF1 has been shown to regulate gene expression during myogenesis [52]. Several myogenic genes (i.e., *myogenin, MyoD*) are known targets of MTF1, which binds their promoter regions during differentiation to activate their expression. The addition of Cu enhances this effect [78]. To determine whether MTF1 binds *mPiC2* during differentiation, previously generated chromatin immunoprecipitation sequencing (ChIP-seq) data for MTF1 binding [78] was accessed and read enrichment over the *PiC2* gene were examined (**Fig. 2B**). MTF1 was enriched at the *PiC2* promoter region (indicated by red box) in cells grown in both the presence and absence of insulin, though the addition of insulin enhanced this effect. MTF1 enrichment was most striking in cells differentiated in the presence of Cu. These results were validated by ChIP-qPCR (**Fig. 2C**). MTF1 enrichment at the *PiC2* promoter was approximately 2-fold higher in cells grown in the presence of insulin compared to those grown in the absence of it. The addition of Cu to medium lacking insulin enhanced MTF1 enrichment 8-fold compared to cells grown in the absence of insulin and Cu and 6-fold compared to cells grown in the presence of insulin and the absence of Cu. Interestingly, the promotor region of *mPiC2* contains three potential metal responsive elements (MRE) for MTF1 characterized by the 5’-TGCRCNC-3’ consensus sequence [87; 88; 89; 90; 91; 92]. The primers used here were designed to cover two of these MREs. To further show that Cu specifically influenced MTF1 binding to the *mPiC2* promoter region, cells grown in the presence of TEPA were analyzed. TEPA treatment decreased MTF1 enrichment markedly, to a level even lower than that observed in cells grown in the absence of both insulin and copper. Cu treatment following TEPA treatment returned MTF1 enrichment levels to those observed in cells grown in the absence of both insulin and Cu. Taken together, these data show that *mPiC2* is a target of MTF1 during primary myoblast differentiation, and that the addition of Cu to the growth medium enhances MTF1 enrichment at the *mPiC2* promoter region.

### Recombinant PiC2 binds Cu^+^ *in vitro*

Sequence analysis of yeast yPiC2 has shown that it is partially homologous to mammalian SLC25A3. The mammalian SLC25A3, or PiC2, has two isoforms: PiC2A and PiC2B. Analysis of the amino acid sequences of mPiC2B and the human hPiC2B (**Fig. 3A**) show that they share a high level of homology (residues highlighted in black). Previous structural characterization of the bovine SLC25A3, bPiC2, suggested the protein has six TM domains [48] and a significant number of conserved Cys, Met, and His residues within these domains, which could indicate possible Cu binding sites. These residues (indicated in **Fig. 3A** with green asterisks; indicated in **Fig. 3B** in green) are also highly conserved in both the human hPiC2B and murine mPiC2B. **Fig. 3B** shows the putative structure of hPiC2B modeled using Alpha fold. Within the TM domain, which is predicted to be comprised of 6 TM segments, are three putative Cu^+^-binding sites. The first possibly consists of Cys90, His79, and Cys75; the second of Cys135, Cys67, and Cys256; and the third of His281, Cys276, and C236. Cu binding experiments using the two purified human PiC2 isoforms (**Fig. 3C**) revealed that PiC2A, had a binding stoichiometry of 2 Cu^+^ per protein, while the other isoform, PiC2B, had a stoichiometry of approximately 3 Cu^+^ per protein. These findings are well in agreement with the Alpha fold prediction that the transporter has 2-3 Cu^+^-binding sites.

**Fig. 3.**
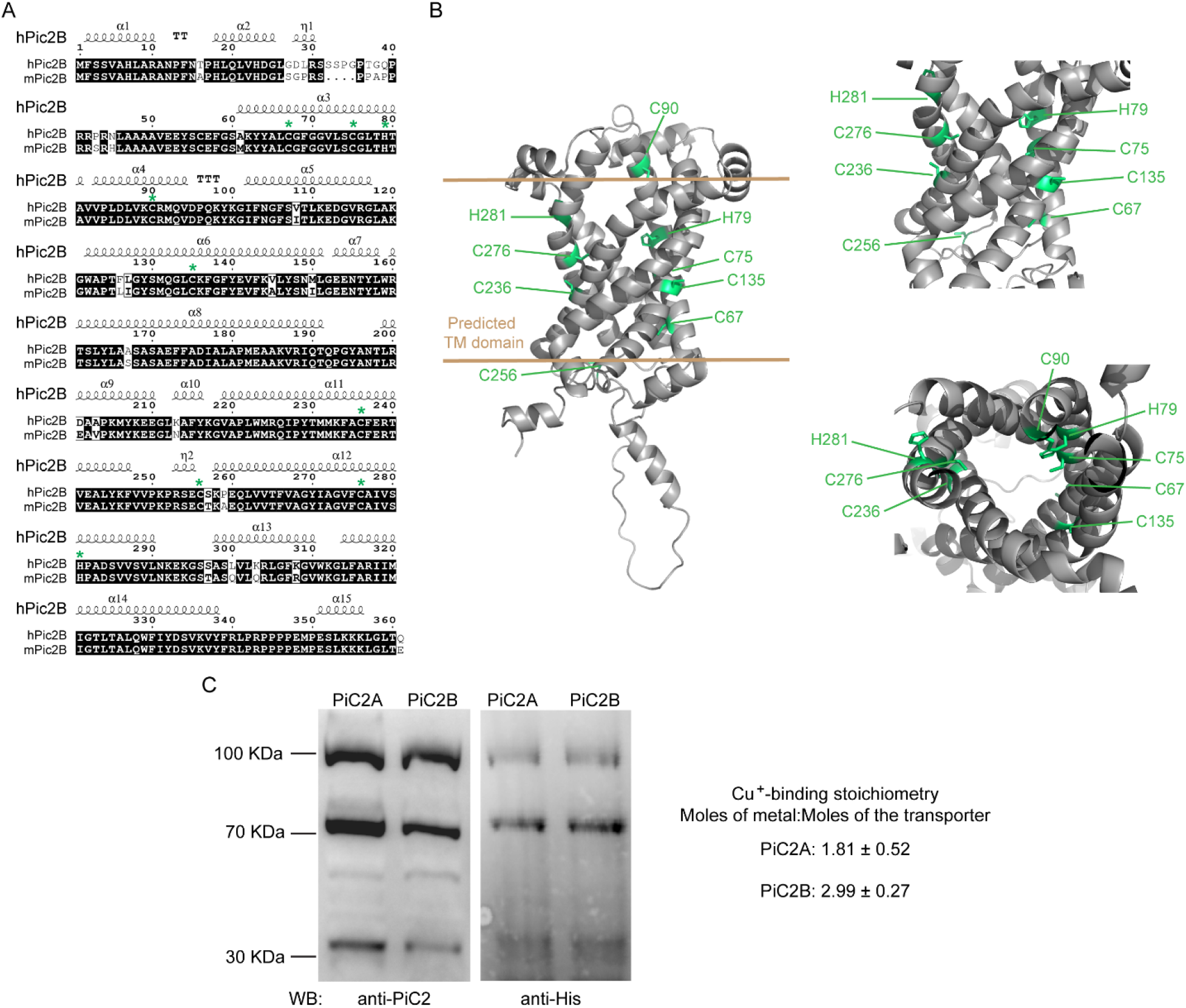
mPiC2 protein has several potential Cu binding sites. **(A)** Sequence homology comparison of the human hPiC2B and murine mPiC2B. Alignment of hPiC2B and mPiC2B amino acid sequences. Areas of homology between the sequences of the two transporters are indicated in black, potential Cu^+^-binding resides indicated with green asterisks (*). **(B)** Representative model of mPic2 protein and potential Cu^+^-binding sites obtained from Alpha fold. Potential Cu^+^-binding residues indicated in green. **(C)** Representative western blots of recombinant PiC2A and PiC2B protein expression detected by both anti-PiC2 antibody and anti-His tag antibody. Monomeric, dimeric, and trimeric forms are detected by both antibodies. Cu^+^-binding stoichiometry of Pic2A and Pic2B. *n*=3, results reported as means ± SEM.

### PiC2 interacts with COX and other mitochondrial cuproproteins in primary myoblasts

COX requires Cu for proper activity and function. Inactivation of PiC2 in both yeast and mice has been shown to result in disruptions of the COX activity and decreases in mitochondrial Cu content [46; 53; 93]. This suggests that PiC2 may interact with COX, or other mitochondrial Cu^+^-binding proteins, such as Sco1 and Sco2, to support the metallation of the oxidase. To test this, the localization of PiC2 relative to the mitochondrial COX in differentiating primary myoblasts was analyzed (**Fig. 4A**). Consistent with the data shown in **Fig. 1A**, mPiC2 staining (red) showed a punctate cytosolic pattern during both proliferation and differentiation. Moreover, the staining pattern of PiC2 was indicative of a mitochondrial distribution of the protein, which was further confirmed by its co-localization with COX (green staining; co-localization indicated by yellow staining). Strikingly, the co-localization of PiC2 with COX appears to be enhanced by the addition of Cu to the growth medium. To further investigate the potential interaction between PiC2 and COX, the proteins were co-immunoprecipitated (co-IP) in proliferating and differentiating myoblasts in the presence and absence of Cu (**Fig. 4B**). As expected, more PiC2 was pulled down in the differentiated samples, consistent with an increase in the expression of PiC2 upon differentiation. Pull down of PiC2 followed by probing for two of the COX subunits, COX1 and COX2, revealed that both subunits are present on the gel, indicating an interaction with PiC2. Moreover, probing for additional mitochondrial cuproproteins Sco1 and Sco2 also resulted in protein signal, showing that these proteins also associate with PiC2 (**Fig. 4B**). These data suggest that PiC2 interacts with at least two types of mitochondrial proteins involved in Cu transport and homeostasis. Interestingly, the interaction between Cu^+^, PiC2 and COX was further confirmed by x-ray fluorescence-mass spectrometry (XRF/MS; **Fig. 4C**). Whole cell extracts of differentiating myoblasts treated with or without insulin or Cu were separated by non-denaturing PAGE to preserve protein-Cu interaction prior to XRF-MS. Samples were analyzed for the presence of Cu, and a high molecular weight band was detected (**Fig. 4C**, indicated by black box). Mass spectrometry sequence analyses showed that COX was the only known cuproprotein identified in the samples, however, PiC2 was also recognized. *In vitro* Cu^+^ transfer experiments using apo-CuA and PiC2A showed that the metallated transporter is capable to donate Cu^+^ to the oxidase, confirming a direct interaction between these proteins.

**Fig. 4.**
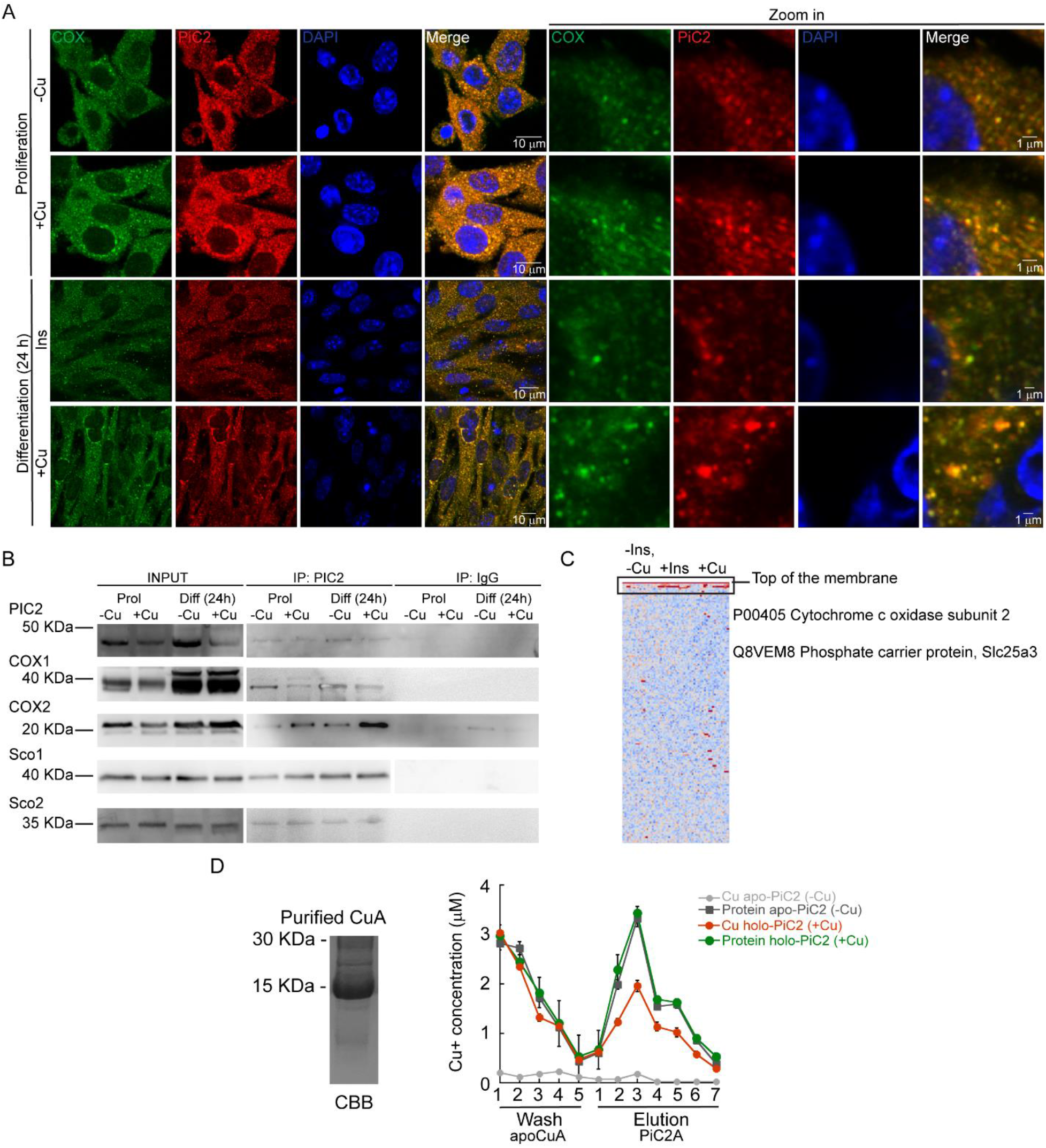
mPiC2 interacts with cytochrome c oxidase and other mitochondrial cuproproteins in primary myoblasts derived from mouse satellite cells. **(A)** mPiC2 co-localizes with COX. Confocal imaging showing mPiC2 (red) and COX (green) co-localization (yellow). The nucleus is stained with DAPI (blue). Set of four panels on the right are zooms of areas in the set of four panels on the left. **(B)** Co-IP of mPiC2 with mitochondrial cuproproteins. Proliferating and differentiating primary myoblasts were treated with or without Cu. Co-IP was performed probing for PiC2 and COX1, COX2, Sco1, or Sco2. **(C)** Synchrotron based X-Ray fluorescence analyses couple to protein sequencing by mass spectrometry of native PAGE gels from whole cell extracts of differentiating primary myoblasts showed the presence of mPiC2 and COX in a high molecular band that contained copper (indicated by black box). **(D)** Cu^+^ transfer from metallated PiC2A to apo-CuA. Cu^+^ concentration is shown in red, and the corresponding protein is shown in green. Control Cu^+^ transfer experiment is shown in pale (Cu^+^ signal) and dark gray (protein signal). In all cases, the contents of the wash and elution fractions are shown. The data corresponds to three independent replicates shown as the mean for Cu^+^ and protein concentration ± SE.

### mPiC2 is required for the growth and differentiation of primary myoblasts derived from mouse satellite cells

Dysfunction and loss of activity of mPiC2, in addition to leading to defects in COX activity, has also been linked to rare diseases, such as fatal and benign infantile myopathies, lactic acidosis, hypertrophic cardiomyopathy, and muscular hypotonia [53; 94]. Mutations in mPiC2 have been shown to lead to disruptions in Pi transport that impacts muscle function and development [94; 95]. To assess the link between mPiC2 function and myogenesis, the CRISPR/Cas9 system was used to knock out (KO) *mPiC2* in primary myoblasts derived from mouse satellite cells. Western blotting (**Fig. 5A**) shows that *mPiC2* KO was successful, with effectively no mPic2 protein signal in the KO cells in both the presence and absence of Cu compared to the control cells transduced with the empty vector. Notably, the *mPiC2* KO had a large impact on the abundance of the mitochondrial cuproproteins, COX1, COX2, Sco1, and Sco2. Addition of Cu to the culture media reverted the impact of this deletion, restoring the protein levels. This effect on COX expression was more pronounced in differentiated cells than in proliferating cells.

**Fig. 5.**
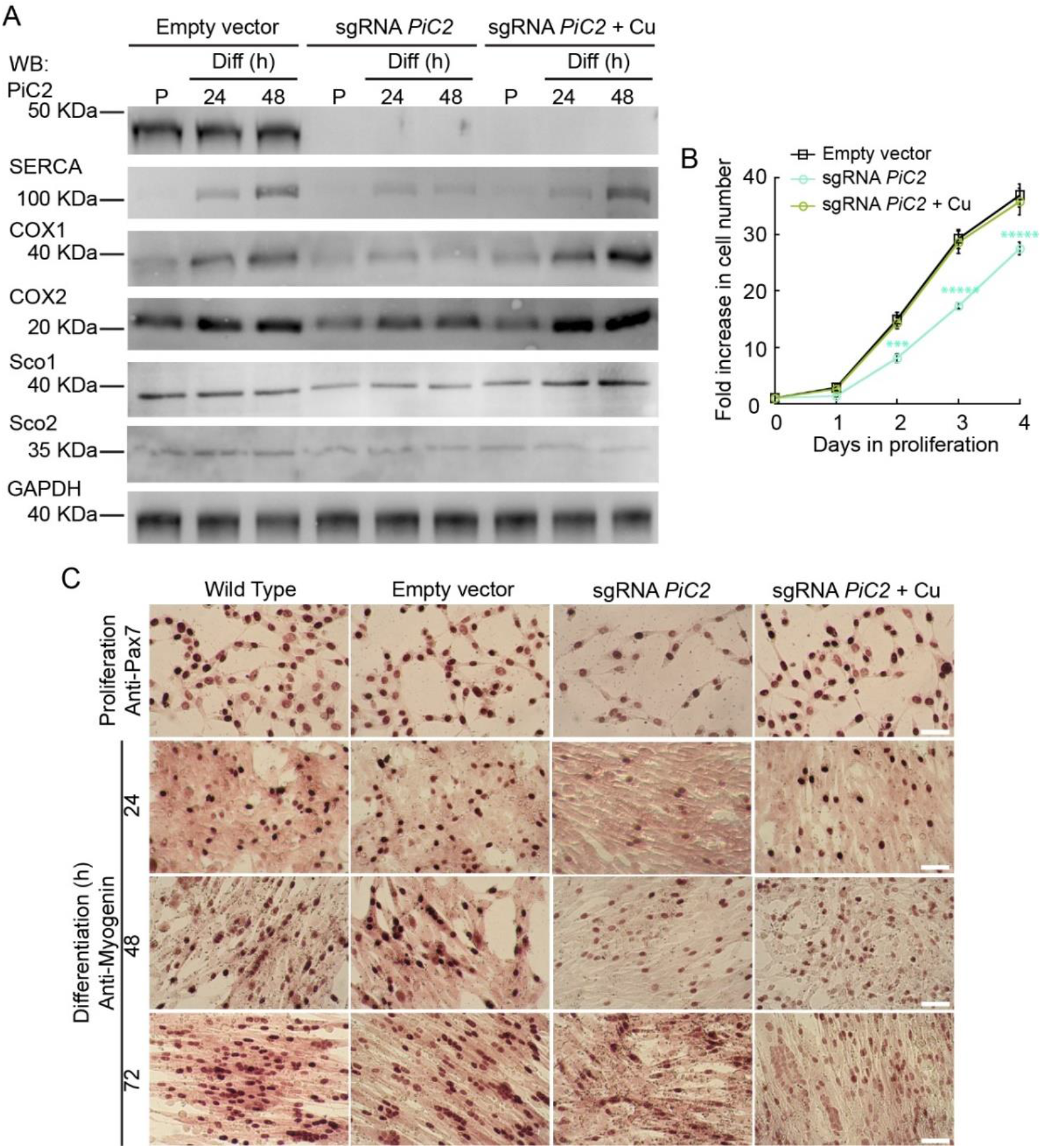
Deletion of *mPiC2* impairs the proliferation and differentiation of primary myoblasts derived from mouse satellite cells. **(A)** Deletion of mPiC2 reduces the expression of mitochondrial cuproproteins. Representative western blot of mPiC2 and mitochondrial cuproproteins in control cells (empty vector) or cells in which CRISPR/Cas9 was used to delete mPiC2 (sgRNA *mPiC2*) or KO cells supplemented with CuSO4 (sgRNA *mPiC2* + Cu) during proliferation (P) and at 24 and 48 h after inducing differentiation. GAPDH was used as a loading control. **(B)** Deletion of mPiC2 delays proliferation. Cell counting assay of empty vector, sgRNA against *mPiC2*, and sgRNA *mPiC2* +Cu cells at different points during proliferation, *n*=3. All values are reported as means ± SE. Significance was determined by two-way ANOVA; *** *p*<0.001 and **** *p*<0.0001 compared to empty vector cells. **(C)** Deletion of mPiC2 delays myogenesis. Staining of wild type, empty vector, sgRNA *mPiC2*, and sgRNA *mPiC2* + Cu cells with either anti-Pax7 antibody (marker of proliferation) or anti-myogenin antibody (marker of differentiation).

The data in **Fig. 1** and **Fig. 2A** showed that mPiC2 expression is upregulated upon differentiation in primary myoblasts and that this effect is enhanced by the addition of Cu to the culture medium. Analysis of the effect that *PiC2* KO has on primary myoblasts revealed that deletion of the transporter impacts also on proliferation, and the cells present a growth defect compared to control cells (**Fig. 5B**). This delay was reversed by the addition of Cu to the growth medium, with the number of cells actively proliferating similarly to control levels. Similarly, immunostaining with the proliferation marker Pax7 (**Fig. 5C**) showed fewer myoblasts in a proliferative state upon *mPiC2* deletion compared to controls, with the presence of Cu mitigating this effect. To determine how *mPiC2* KO affects myogenesis, cells were stained with an anti-myogenin hybridoma, a marker of differentiation. As with proliferation, *mPiC2* KO resulted in a delay in the progress of differentiation compared to the control cells, an effect that the addition of Cu to the growth medium abolished. Taken together, the experiments presented here confirmed that mPiC2 is a mitochondrial Cu transporter that interacts with additional cuproproteins to favor maturation of COX. This process favors the development and differentiation of cultured skeletal muscle cells. However, despite that mPiC2 seems to contribute to this process, it is not an essential protein for muscle maturation and additional mechanisms are in place, such as COX17, and others.

## DISCUSSION

Mitochondrial function and, subsequently, cellular energy production are reliant in part on Cu. While Cu transport within the mitochondria has been characterized, the delivery of Cu to the mitochondria through the Cu-impermeable mitochondrial membranes is poorly understood. Studies in yeast identified a mitochondrial phosphate transporter, yPiC2, that is necessary for the import of Cu into the mitochondria and the metallation of COX [46]. The closest mammalian homolog of this transporter, SLC25A3 (PiC2), has mainly been characterized as a mitochondrial transporter of Pi. Our studies in murine primary myoblasts derived from mouse satellite cells, an ideal system for studying mitochondrial biogenesis and transport, have revealed a role for PiC2 in this system.

Muscle tissue has an abundance of mitochondria, as these tissues have high energy requirements. This subsequently leads to a high demand in Cu to metalate cuproproteins involved in energy production and mitochondrial function, such as COX [46]. To meet these needs, myoblasts must mobilize additional Cu to the mitochondria. We found that mPiC2 expression is induced in primary myoblasts upon differentiation. This finding suggests a role for PiC2 in myogenesis, though whether this role was related to Cu delivery or not was yet unclear.

In addition to being required for the induction of myogenesis, Cu is also implicated in the transcriptional regulation of myogenic genes [52]. Myoblast differentiation is induced *in vitro* by serum starvation and supplementation of insulin into the culture medium, and the lack of insulin in the medium results in cells that differentiate poorly. We show that while insulin depletion impedes the upregulation of *PiC2* in differentiating myoblasts, the addition of Cu to the growth medium restores the expression of the transporter to that observed in cells supplemented in insulin. Conversely, the depletion of Cu via the chelator TEPA resulted in *mPiC2* expression levels on par with those observed in insulin-deprived myoblasts, indicating cells without Cu differentiate poorly. Adding Cu back following chelation with TEPA reverses this effect. These data together imply that Cu is both necessary for the differentiation-induced increase in *mPiC2* expression and sufficient to ameliorate the effects of insulin-deprivation on the differentiated myoblasts.

We have previously described the metal-sensing transcription factor MTF1 as playing a key role in myoblast differentiation. MTF1 binds the promoter regions of myogenic genes to induce their expression, and this event is enhanced by the addition of Cu to the myoblast growth medium [78]. Similarly, we observed a marked enrichment in MTF1 binding to the promoter region of the *mPiC2* gene during myogenesis in the presence of Cu. This enrichment is even several-fold higher than that observed in differentiating myoblasts induced by the addition of insulin, suggesting the presence of Cu additionally enhances the *MTF1-mPiC2* promoter binding. Depletion of Cu greatly impedes the binding of MTF1 to the promoter of the transporter, and the supplementation of Cu to the medium following Cu depletion restores partially the binding to roughly 50% of that seen in insulin-induced cells. These data suggest that enhanced MTF1 binding to the *mPiC2* promoter during differentiation could cause the RNA- and protein-level increases in mPiC2 expression observed in differentiated cells and that binding of MTF1 to the promoter region is promoted by the addition of Cu. In this way, the overall cellular increase in Cu can facilitate the delivery of Cu to specific compartments, such as the mitochondria, to be loaded into proteins that require Cu as a cofactor for their enzymatic activity.

For PiC2 to act as a Cu^+^-transporter, the protein needs to contain residues that coordinate Cu binding, such as Cys, His, and Met. yPiC2, the first SLC25A3 to be identified as a Cu transporter, has several such residues, many of which are conserved in the mammalian homologs. Sequence analysis between the human and mouse B isoforms of PiC2 (hPiC2B and mPiC2B, respectively) revealed conserved residues and domains that are common to other Cu transporters. Homology modeling of mPiC2B revealed three potential metal binding sites, which were validated by biochemical analyses showed that PiC2 is, indeed, a Cu^+^-binding protein.

The current understanding of COX metallation involves the soluble protein, Cox17, and two proteins located in the inner membrane of the mitochondria, Sco1 and Sco2 [96; 97; 98; 99; 100]. As of yet, there has been no mechanism describing how Cu gets in through the outer mitochondrial membrane to these proteins in order to metalate COX. Based on the fundamental principle that there is no labile Cu in cells due to its reactivity, it is therefore necessary that Cu is transported from donor to acceptor proteins via direct interaction. It is tempting to speculate that PiC2 might fit this role. Confocal imaging of immunostained PiC2 and COX1 revealed that the two co-localize in a punctate pattern indicative of mitochondrial localization. Co-IP and XRF/MS analyses further confirmed that there might be a functional PiC2-COX interaction within differentiating primary myoblasts. PiC2 interacting directly with both COX1 and COX2 subunits— presumably to transfer Cu to COX—is especially interesting, as that would suggest some redundancy with Cox17, Sco1, and Sco2, to ensure COX metallation, and activation to successfully perform respiration processes. Whether or not mitochondrial Cu^+^ delivery via PiC2 is in any way coupled to mitochondrial Pi transport is unknown and requires additional study.

mPiC2 expression was significantly upregulated during myoblast differentiation, which suggests a requirement for the transporter during myogenesis but does not directly prove it. CRISPR/Cas9-mediated deletion of *mPiC2* in myoblasts resulted in the reduction in the expression of COX1, COX2, Sco1, and Sco2, as well as the delayed entry of the cells into both proliferation and differentiation states. These data indicates that mPic2 expression contributes to normal myoblast growth and differentiation. Interestingly, the supplementation of Cu to the culture medium of *mPiC2* KO myoblasts completely reversed these effects. This strongly suggests that the driving force behind the deleterious effects of *mPiC2* KO is the lack of Cu delivery to mitochondria. However, this does imply that Cu may still be getting into the impermeable outer mitochondrial membrane even in the absence of mPiC2. This phenomenon will require further investigation.

Taken together, the data demonstrate that mPiC2 expression is required both for the proper growth of proliferating myoblasts and the progression of myogenesis. *mPiC2* expression is upregulated during the differentiation of primary myoblasts, likely enhanced by the binding of MTF1 to the promoter region of the *mPiC2* gene. The transporter also interacts with mitochondrial cuproproteins, such as COX and Sco1/2, suggesting a possible role for mPiC2 in metalating these proteins. Deletion of *PiC2* results in the downregulation of mitochondrial cuproproteins, such as COX, as well as the delay of both proliferation and differentiation. The addition of Cu to the growth medium is sufficient to offset the effects of *mPiC2* deletion, suggesting that the transporter plays a role in the network of Cu homeostasis in myoblasts. This study provides a foundation for additional study of mitochondrial Cu delivery and the role that Cu plays in both mitochondrial biogenesis and myogenesis in muscle tissue.

## AUTHOR CONTRIBUTIONS

T.P.-B. conceived and designed the research; C.M.C., M.Q., I.A., M.N.M., A.I.C., E.J.D. A.L. and T.P.-B. performed experiments and compiled data; C.M.C., A.L., A.J.V., P.C., J.G.N. and T.P.-B. analyzed data; C.M.C. and T.P.-B. prepared figures and tables; C.M.C and T.P.-B. drafted the manuscript; all authors edited and revised the manuscript; all authors approved the final version of the manuscript.

## ACKNOWLEDGEMENTS

This work was funded by NIH grant R01AR077578 (NIAMS) to T.P.-B. Portions of this work were performed at GeoSoilEnviroCARS (The University of Chicago, Sector 13), Advanced Photon Source (APS), Argonne National Laboratory. GeoSoilEnviroCARS is supported by the National Science Foundation - Earth Sciences (EAR - 1634415). This research used resources of the Advanced Photon Source; a U.S. Department of Energy (DOE) Office of Science User Facility operated for the DOE Office of Science by Argonne National Laboratory under Contract No. DE-AC02-06CH11357. M.Q. was recipient of the 2022 ASBMB Undergraduate Research Award.

The authors are thankful to Dr. Pablo Reyes-Gutierrez for his technical support and to Drs. Paul Cobine, Monserrat Olea-Flores and Ms. Ella Y. Kim for their critical comments on this manuscript.

## REFERENCES

[1] J.J.R. Fraústo da Silva, and R.J.P. Williams, The biological chemistry of the elements: the inorganic chemistry of life, Oxford University Press, Oxford, 2001.

[2] M.C. Linder, and M. Hazegh-Azam, Copper biochemistry and molecular biology. The American journal of clinical nutrition 63 (1996) 797S–811S.

[3] J. Prousek, Fenton chemistry in biology and medicine. 79 (2007) 2325–2338.

[4] J.M. Argüello, D. Raimunda, and T. Padilla-Benavides, Mechanisms of copper homeostasis in bacteria. Front Cell Infect Microbiol doi: 10.3389/fcimb.2013.00073 (2013).

[5] N.J. Robinson, and D.R. Winge, Copper metallochaperones. Annu Rev Biochem 79 (2010) 537–62.

[6] R.A. Festa, and D.J. Thiele, Copper: an essential metal in biology. Current biology: CB 21 (2011) R877–83.

[7] H. Ohrvik, and D.J. Thiele, How copper traverses cellular membranes through the mammalian copper transporter 1, Ctr1. Annals of the New York Academy of Sciences 1314 (2014) 32–41.

[8] S. Lutsenko, N.L. Barnes, M.Y. Bartee, and O.Y. Dmitriev, Function and regulation of human copper-transporting ATPases. Physiol Rev 87 (2007) 1011–46.

[9] T.V. O’Halloran, and V.C. Culotta, Metallochaperones, an intracellular shuttle service for metal ions. J Biol Chem 275 (2000) 25057–60.

[10] T.D. Rae, P.J. Schmidt, R.A. Pufahl, V.C. Culotta, and T.V. O’Halloran, Undetectable intracellular free copper: the requirement of a copper chaperone for superoxide dismutase. Science 284 (1999) 805–808.

[11] K.J. Waldron, J.C. Rutherford, D. Ford, and N.J. Robinson, Metalloproteins and metal sensing. Nature 460 (2009) 823–830.

[12] A.N. Barry, A. Otoikhian, S. Bhatt, U. Shinde, R. Tsivkovskii, N.J. Blackburn, and S. Lutsenko, The Lumenal Loop Met(672)-Pro(707) of Copper-transporting ATPase ATP7A Binds Metals and Facilitates Copper Release from the Intramembrane Sites. Journal of Biological Chemistry 286 (2011) 26585–26594.

[13] A. Otoikhian, A.N. Barry, M. Mayfield, M. Nilges, Y.P. Huang, S. Lutsenko, and N.J. Blackburn, Lumenal loop M672-P707 of the Menkes protein (ATP7A) transfers copper to peptidylglycine monooxygenase. J Am Chem Soc 134 (2012) 10458–10468.

[14] T. Padilla-Benavides, C.J. McCann, and J.M. Argüello, The mechanism of Cu^+^ transport ATPases: Interaction with Cu^+^ chaperones and the role of transient metal-binding sites. J Biol Chem 288 (2013) 69–78.

[15] T. Padilla-Benavides, A.M. George Thompson, M.M. McEvoy, and J.M. Arguello, Mechanism of ATPase-mediated Cu+ export and delivery to periplasmic chaperones: the interaction of Escherichia coli CopA and CusF. J Biol Chem 289 (2014) 20492–501.

[16] J. Lee, M.J. Petris, and D.J. Thiele, Characterization of mouse embryonic cells deficient in the ctr1 high affinity copper transporter. Identification of a Ctr1-independent copper transport system. J Biol Chem 277 (2002) 40253–9.

[17] E.B. Maryon, S.A. Molloy, and J.H. Kaplan, Cellular glutathione plays a key role in copper uptake mediated by human copper transporter 1. American journal of physiology. Cell physiology 304 (2013) C768–79.

[18] S. Lutsenko, Copper trafficking to the secretory pathway. Metallomics (2016).

[19] A. Mauro, Satellite cell of skeletal muscle fibers. The Journal of biophysical and biochemical cytology 9 (1961) 493–5.

[20] H. Yin, F. Price, and M.A. Rudnicki, Satellite cells and the muscle stem cell niche. Physiol Rev 93 (2013) 23–67.

[21] A.S. Brack, and T.A. Rando, Tissue-specific stem cells: lessons from the skeletal muscle satellite cell. Cell stem cell 10 (2012) 504–14.

[22] D. Montarras, A. L’Honore, and M. Buckingham, Lying low but ready for action: the quiescent muscle satellite cell. The FEBS journal 280 (2013) 4036–50.

[23] M. Buckingham, and P.W. Rigby, Gene regulatory networks and transcriptional mechanisms that control myogenesis. Developmental cell 28 (2014) 225–38.

[24] N.C. Chang, and M.A. Rudnicki, Satellite cells: the architects of skeletal muscle. Current topics in developmental biology 107 (2014) 161–81.

[25] T. Padilla-Benavides, B.T. Nasipak, and A.N. Imbalzano, Brg1 Controls the Expression of Pax7 to Promote Viability and Proliferation of Mouse Primary Myoblasts. Journal of cellular physiology 230 (2015) 2990–7.

[26] N. Motohashi, and A. Asakura, Muscle satellite cell heterogeneity and self-renewal. Frontiers in cell and developmental biology 2 (2014) 1.

[27] R. Sambasivan, and S. Tajbakhsh, Adult skeletal muscle stem cells. Results and problems in cell differentiation 56 (2015) 191–213.

[28] P. Seale, J. Ishibashi, A. Scime, and M.A. Rudnicki, Pax7 is necessary and sufficient for the myogenic specification of CD45+:Sca1+ stem cells from injured muscle. PLoS biology 2 (2004) E130.

[29] P. Seale, L.A. Sabourin, A. Girgis-Gabardo, A. Mansouri, P. Gruss, and M.A. Rudnicki, Pax7 is required for the specification of myogenic satellite cells. Cell 102 (2000) 777–86.

[30] J. von Maltzahn, A.E. Jones, R.J. Parks, and M.A. Rudnicki, Pax7 is critical for the normal function of satellite cells in adult skeletal muscle. Proc Natl Acad Sci U S A 110 (2013) 16474–9.

[31] Z. Yablonka-Reuveni, M.A. Rudnicki, A.J. Rivera, M. Primig, J.E. Anderson, and P. Natanson, The transition from proliferation to differentiation is delayed in satellite cells from mice lacking MyoD. Developmental biology 210 (1999) 440–55.

[32] J. Huijbregts, J.D. White, and M.D. Grounds, The absence of MyoD in regenerating skeletal muscle affects the expression pattern of basement membrane, interstitial matrix and integrin molecules that is consistent with delayed myotube formation. Acta histochemica 103 (2001) 379–96.

[33] L.A. Megeney, B. Kablar, K. Garrett, J.E. Anderson, and M.A. Rudnicki, MyoD is required for myogenic stem cell function in adult skeletal muscle. Genes & development 10 (1996) 1173–83.

[34] B. Gayraud-Morel, F. Chretien, P. Flamant, D. Gomes, P.S. Zammit, and S. Tajbakhsh, A role for the myogenic determination gene Myf5 in adult regenerative myogenesis. Developmental biology 312 (2007) 13–28.

[35] R.N. Cooper, S. Tajbakhsh, V. Mouly, G. Cossu, M. Buckingham, and G.S. Butler-Browne, In vivo satellite cell activation via Myf5 and MyoD in regenerating mouse skeletal muscle. Journal of cell science 112 (Pt 17) (1999) 2895–901.

[36] P.S. Zammit, J.P. Golding, Y. Nagata, V. Hudon, T.A. Partridge, and J.R. Beauchamp, Muscle satellite cells adopt divergent fates: a mechanism for self-renewal? The Journal of cell biology 166 (2004) 347–57.

[37] C.D. Moyes, O.A. Mathieu-Costello, N. Tsuchiya, C. Filburn, and R.G. Hansford, Mitochondrial biogenesis during cellular differentiation. The American journal of physiology 272 (1997) C1345–51.

[38] A.H. Remels, R.C. Langen, P. Schrauwen, G. Schaart, A.M. Schols, and H.R. Gosker, Regulation of mitochondrial biogenesis during myogenesis. Molecular and cellular endocrinology 315 (2010) 113–20.

[39] N.H. Herzberg, R. Zwart, R.A. Wolterman, J.P. Ruiter, R.J. Wanders, P.A. Bolhuis, and C. van den Bogert, Differentiation and proliferation of respiration-deficient human myoblasts. Bba-Bioenergetics 1181 (1993) 63–7.

[40] N. Hamai, M. Nakamura, and A. Asano, Inhibition of mitochondrial protein synthesis impaired C2C12 myoblast differentiation. Cell structure and function 22 (1997) 421–31.

[41] S.C. Leary, B.J. Battersby, R.G. Hansford, and C.D. Moyes, Interactions between bioenergetics and mitochondrial biogenesis. Bba-Bioenergetics 1365 (1998) 522–30.

[42] S.C. Leary, B.C. Hill, C.N. Lyons, C.G. Carlson, D. Michaud, C.S. Kraft, K. Ko, D.M. Glerum, and C.D. Moyes, Chronic treatment with azide in situ leads to an irreversible loss of cytochrome c oxidase activity via holoenzyme dissociation. J Biol Chem 277 (2002) 11321–8.

[43] A. Wagatsuma, and K. Sakuma, Mitochondria as a potential regulator of myogenesis. TheScientificWorldJournal 2013 (2013) 593267.

[44] S.C. Leary, B.A. Kaufman, G. Pellecchia, G.H. Guercin, A. Mattman, M. Jaksch, and E.A. Shoubridge, Human SCO1 and SCO2 have independent, cooperative functions in copper delivery to cytochrome c oxidase. Human molecular genetics 13 (2004) 1839–48.

[45] M.N. Morgada, L.A. Abriata, C. Cefaro, K. Gajda, L. Banci, and A.J. Vila, Loop recognition and copper-mediated disulfide reduction underpin metal site assembly of CuA in human cytochrome oxidase. Proc Natl Acad Sci U S A 112 (2015) 11771–6.

[46] K.E. Vest, S.C. Leary, D.R. Winge, and P.A. Cobine, Copper import into the mitochondrial matrix in Saccharomyces cerevisiae is mediated by Pic2, a mitochondrial carrier family protein. J Biol Chem 288 (2013) 23884–92.

[47] E.L. Seifert, E. Ligeti, J.A. Mayr, N. Sondheimer, and G. Hajnoczky, The mitochondrial phosphate carrier: Role in oxidative metabolism, calcium handling and mitochondrial disease. Biochemical and biophysical research communications 464 (2015) 369–75.

[48] M.J. Runswick, S.J. Powell, P. Nyren, and J.E. Walker, Sequence of the bovine mitochondrial phosphate carrier protein: structural relationship to ADP/ATP translocase and the brown fat mitochondria uncoupling protein. The EMBO journal 6 (1987) 1367–73.

[49] V. Dolce, V. Iacobazzi, F. Palmieri, and J.E. Walker, The sequences of human and bovine genes of the phosphate carrier from mitochondria contain evidence of alternatively spliced forms. J Biol Chem 269 (1994) 10451–60.

[50] V. Dolce, G. Fiermonte, and F. Palmieri, Tissue-specific expression of the two isoforms of the mitochondrial phosphate carrier in bovine tissues. FEBS letters 399 (1996) 95–8.

[51] G. Fiermonte, V. Dolce, and F. Palmieri, Expression in Escherichia coli, functional characterization, and tissue distribution of isoforms A and B of the phosphate carrier from bovine mitochondria. J Biol Chem 273 (1998) 22782–7.

[52] C. Tavera-Montanez, S.J. Hainer, D. Cangussu, S.J.V. Gordon, Y. Xiao, P. Reyes-Gutierrez, A.N. Imbalzano, J.G. Navea, T.G. Fazzio, and T. Padilla-Benavides, The classic metal-sensing transcription factor MTF1 promotes myogenesis in response to copper. Faseb J 33 (2019) 14556–14574.

[53] A. Boulet, K.E. Vest, M.K. Maynard, M.G. Gammon, A.C. Russell, A.T. Mathews, S.E. Cole, X. Zhu, C.B. Phillips, J.Q. Kwong, S.C. Dodani, S.C. Leary, and P.A. Cobine, The mammalian phosphate carrier SLC25A3 is a mitochondrial copper transporter required for cytochrome c oxidase biogenesis. J Biol Chem 293 (2018) 1887–1896.

[54] B.T. Nasipak, T. Padilla-Benavides, K.M. Green, J.D. Leszyk, W. Mao, S. Konda, S. Sif, S.A. Shaffer, Y. Ohkawa, and A.N. Imbalzano, Opposing calcium-dependent signalling pathways control skeletal muscle differentiation by regulating a chromatin remodelling enzyme. Nature communications 6 (2015) 7441.

[55] K.E. Vest, A.L. Paskavitz, J.B. Lee, and T. Padilla-Benavides, Dynamic changes in copper homeostasis and post-transcriptional regulation of Atp7a during myogenic differentiation. Metallomics 10 (2018) 309–322.

[56] H.L. Goel, and C.S. Dey, Focal adhesion kinase tyrosine phosphorylation is associated with myogenesis and modulated by insulin. Cell proliferation 35 (2002) 131–42.

[57] R. Conejo, and M. Lorenzo, Insulin signaling leading to proliferation, survival, and membrane ruffling in C2C12 myoblasts. Journal of cellular physiology 187 (2001) 96–108.

[58] N.E. Sanjana, O. Shalem, and F. Zhang, Improved vectors and genome-wide libraries for CRISPR screening. Nature methods 11 (2014) 783–784.

[59] O. Shalem, N.E. Sanjana, E. Hartenian, X. Shi, D.A. Scott, T. Mikkelson, D. Heckl, B.L. Ebert, D.E. Root, J.G. Doench, and F. Zhang, Genome-scale CRISPR-Cas9 knockout screening in human cells. Science 343 (2014) 84–87.

[60] P. Sheffield, S. Garrard, and Z. Derewenda, Overcoming expression and purification problems of RhoGDI using a family of “parallel” expression vectors. Protein Expr Purif 15 (1999) 34–9.

[61] K.J. Livak, and T.D. Schmittgen, Analysis of relative gene expression data using real-time quantitative PCR and the 2(-Delta Delta C(T)) Method. Methods 25 (2001) 402–8.

[62] M.M. Bradford, A rapid and sensitive method for the quantitation of microgram quantities of protein utilizing the principle of protein-dye binding. Anal Biochem 72 (1976) 248–54.

[63] R.C. Edgar, MUSCLE: multiple sequence alignment with high accuracy and high throughput. Nucleic Acids Res 32 (2004) 1792–7.

[64] P. Gouet, E. Courcelle, D.I. Stuart, and F. Metoz, ESPript: analysis of multiple sequence alignments in PostScript. Bioinformatics 15 (1999) 305–8.

[65] X. Robert, and P. Gouet, Deciphering key features in protein structures with the new ENDscript server. Nucleic Acids Res 42 (2014) W320–4.

[66] J. Jumper, R. Evans, A. Pritzel, T. Green, M. Figurnov, O. Ronneberger, K. Tunyasuvunakool, R. Bates, A. Zidek, A. Potapenko, A. Bridgland, C. Meyer, S.A.A. Kohl, A.J. Ballard, A. Cowie, B. Romera-Paredes, S. Nikolov, R. Jain, J. Adler, T. Back, S. Petersen, D. Reiman, E. Clancy, M. Zielinski, M. Steinegger, M. Pacholska, T. Berghammer, S. Bodenstein, D. Silver, O. Vinyals, A.W. Senior, K. Kavukcuoglu, P. Kohli, and D. Hassabis, Highly accurate protein structure prediction with AlphaFold. Nature 596 (2021) 583–589.

[67] M. Varadi, S. Anyango, M. Deshpande, S. Nair, C. Natassia, G. Yordanova, D. Yuan, O. Stroe, G. Wood, A. Laydon, A. Zidek, T. Green, K. Tunyasuvunakool, S. Petersen, J. Jumper, E. Clancy, R. Green, A. Vora, M. Lutfi, M. Figurnov, A. Cowie, N. Hobbs, P. Kohli, G. Kleywegt, E. Birney, D. Hassabis, and S. Velankar, AlphaFold Protein Structure Database: massively expanding the structural coverage of protein-sequence space with high-accuracy models. Nucleic Acids Res 50 (2022) D439–D444.

[68] F.W. Studier, Protein production by auto-induction in high density shaking cultures. Protein Expr Purif 41 (2005) 207–34.

[69] M. Gonzalez-Guerrero, and J.M. Arguello, Mechanism of Cu+-transporting ATPases: soluble Cu+ chaperones directly transfer Cu+ to transmembrane transport sites. Proc Natl Acad Sci U S A 105 (2008) 5992–7.

[70] A.K. Mandal, and J.M. Argüello, Functional roles of metal binding domains of the *Archaeoglobus fulgidus* Cu^+^-ATPase CopA. Biochemistry 42 (2003) 11040–11047.

[71] D. Raimunda, J.E. Long, T. Padilla-Benavides, C.M. Sassetti, and J.M. Arguello, Differential roles for the Co(2+) /Ni(2+) transporting ATPases, CtpD and CtpJ, in Mycobacterium tuberculosis virulence. Mol Microbiol 91 (2014) 185–97.

[72] C.E. Blaby-Haas, T. Padilla-Benavides, R. Stube, J.M. Arguello, and S.S. Merchant, Evolution of a plant-specific copper chaperone family for chloroplast copper homeostasis. Proc Natl Acad Sci U S A 111 (2014) E5480–7.

[73] T. Padilla-Benavides, J.E. Long, D. Raimunda, C.M. Sassetti, and J.M. Argüello, A novel P1B-type Mn^2+^-transporting ATPase is required for secreted protein metallation in mycobacteria. J Biol Chem 288 (2013) 11334–11347.

[74] A.J. Leguto, M.A. Smith, M.N. Morgada, U.A. Zitare, D.H. Murgida, K.M. Lancaster, and A.J. Vila, Dramatic Electronic Perturbations of CuA Centers via Subtle Geometric Changes. J Am Chem Soc 141 (2019) 1373–1381.

[75] T. Padilla-Benavides, and J.M. Arguello, Assay of Copper Transfer and Binding to P1B-ATPases. Methods Mol Biol 1377 (2016) 267–77.

[76] M. Olea-Flores, J. Kan, A. Carlson, S.A. Syed, C. McCann, V. Mondal, C. Szady, H.M. Ricker, A. McQueen, J.G. Navea, L.A. Caromile, and T. Padilla-Benavides, ZIP11 Regulates Nuclear Zinc Homeostasis in HeLa Cells and Is Required for Proliferation and Establishment of the Carcinogenic Phenotype. Front Cell Dev Biol 10 (2022) 895433.

[77] C.A. Vilchis-Nestor, M.L. Roldan, A. Leonardi, J.G. Navea, T. Padilla-Benavides, and L. Shoshani, Ouabain Enhances Cell-Cell Adhesion Mediated by beta1 Subunits of the Na(+),K(+)-ATPase in CHO Fibroblasts. International journal of molecular sciences 20 (2019).

[78] C. Tavera-Montanez, S.J. Hainer, D. Cangussu, S.J.V. Gordon, Y. Xiao, P. Reyes-Gutierrez, A.N. Imbalzano, J.G. Navea, T.G. Fazzio, and T. Padilla-Benavides, The classic metal-sensing transcription factor MTF1 promotes myogenesis in response to copper. FASEB J 33 (2019) 14556–14574.

[79] S.J.V. Gordon, Y. Xiao, A.L. Paskavitz, N. Navarro-Tito, J.G. Navea, and T. Padilla-Benavides, Atomic Absorbance Spectroscopy to Measure Intracellular Zinc Pools in Mammalian Cells. Journal of visualized experiments: JoVE (2019).

[80] A.L. Paskavitz, J. Quintana, D. Cangussu, C. Tavera-Montanez, Y. Xiao, S. Ortiz-Miranda, J.G. Navea, and T. Padilla-Benavides, Differential expression of zinc transporters accompanies the differentiation of C2C12 myoblasts. J Trace Elem Med Biol 49 (2018) 27–34.

[81] T. Padilla-Benavides, M.L. Roldan, I. Larre, D. Flores-Benitez, N. Villegas-Sepulveda, R.G. Contreras, M. Cereijido, and L. Shoshani, The polarized distribution of Na+,K+-ATPase: role of the interaction between {beta} subunits. Molecular biology of the cell 21 (2010) 2217–25.

[82] D. Raimunda, T. Khare, C. Giometti, S. Vogt, J.M. Argüello, and L. Finney, Identifying metalloproteins through X-ray fluorescence mapping and mass spectrometry. Metallomics 4 (2012) 921–927.

[83] D. Raimunda, T. Padilla-Benavides, S. Vogt, S. Boutigny, K.N. Tomkinson, L.A. Finney, and J.M. Argüello, Periplasmic response upon disruption of transmembrane Cu^+^ transport in *Pseudomonas aeruginosa*. Metallomics 5 (2013) 144–151.

[84] S. Sutton, Lanzirotti, A., Newville, M., Rivers, M., Eng, P. Lefticariu, L, Spatially-Resolved Elemental Analysis, Spectroscopy and Diffraction at the GSECARS Sector at the Advanced Photon Source. Journal of Environmental Quality 46 (2017) 1158–1165.

[85] M. Newville, Larch: an analysis package for XAFS and related spectroscopies.. Journal of Physics: Conference Series. IOP Publishing. 430 (2013) 012007.

[86] R. Conejo, A.M. Valverde, M. Benito, and M. Lorenzo, Insulin produces myogenesis in C2C12 myoblasts by induction of NF-kappaB and downregulation of AP-1 activities. Journal of cellular physiology 186 (2001) 82–94.

[87] Y. Wang, I. Lorenzi, O. Georgiev, and W. Schaffner, Metal-responsive transcription factor-1 (MTF-1) selects different types of metal response elements at low vs. high zinc concentration. Biological chemistry 385 (2004) 623–32.

[88] A. Selvaraj, K. Balamurugan, H. Yepiskoposyan, H. Zhou, D. Egli, O. Georgiev, D.J. Thiele, and W. Schaffner, Metal-responsive transcription factor (MTF-1) handles both extremes, copper load and copper starvation, by activating different genes. Genes & development 19 (2005) 891–6.

[89] R. Heuchel, F. Radtke, O. Georgiev, G. Stark, M. Aguet, and W. Schaffner, The transcription factor MTF-1 is essential for basal and heavy metal-induced metallothionein gene expression. The EMBO journal 13 (1994) 2870–5.

[90] S.J. Langmade, R. Ravindra, P.J. Daniels, and G.K. Andrews, The transcription factor MTF-1 mediates metal regulation of the mouse ZnT1 gene. J Biol Chem 275 (2000) 34803–9.

[91] G.K. Andrews, D.K. Lee, R. Ravindra, P. Lichtlen, M. Sirito, M. Sawadogo, and W. Schaffner, The transcription factors MTF-1 and USF1 cooperate to regulate mouse metallothionein-I expression in response to the essential metal zinc in visceral endoderm cells during early development. The EMBO journal 20 (2001) 1114–22.

[92] A.T. Miles, G.M. Hawksworth, J.H. Beattie, and V. Rodilla, Induction, regulation, degradation, and biological significance of mammalian metallothioneins. Critical reviews in biochemistry and molecular biology 35 (2000) 35–70.

[93] X. Zhu, A. Boulet, K.M. Buckley, C.B. Phillips, M.G. Gammon, L.E. Oldfather, S.A. Moore, S.C. Leary, and P.A. Cobine, Mitochondrial copper and phosphate transporter specificity was defined early in the evolution of eukaryotes. eLife 10 (2021).

[94] J.A. Mayr, F.A. Zimmermann, R. Horvath, H.C. Schneider, B. Schoser, E. Holinski-Feder, B. Czermin, P. Freisinger, and W. Sperl, Deficiency of the mitochondrial phosphate carrier presenting as myopathy and cardiomyopathy in a family with three affected children. Neuromuscular disorders: NMD 21 (2011) 803–8.

[95] J.A. Mayr, O. Merkel, S.D. Kohlwein, B.R. Gebhardt, H. Bohles, U. Fotschl, J. Koch, M. Jaksch, H. Lochmuller, R. Horvath, P. Freisinger, and W. Sperl, Mitochondrial phosphate-carrier deficiency: a novel disorder of oxidative phosphorylation. American journal of human genetics 80 (2007) 478–84.

[96] S.C. Leary, F. Sasarman, T. Nishimura, and E.A. Shoubridge, Human SCO2 is required for the synthesis of CO II and as a thiol-disulphide oxidoreductase for SCO1. Human molecular genetics 18 (2009) 2230–40.

[97] L. Banci, I. Bertini, S. Ciofi-Baffoni, A. Janicka, M. Martinelli, H. Kozlowski, and P. Palumaa, A structural-dynamical characterization of human Cox17. J Biol Chem 283 (2008) 7912–20.

[98] L. Banci, I. Bertini, S. Ciofi-Baffoni, T. Hadjiloi, M. Martinelli, and P. Palumaa, Mitochondrial copper(I) transfer from Cox17 to Sco1 is coupled to electron transfer. Proc Natl Acad Sci U S A 105 (2008) 6803–8.

[99] Y.C. Horng, S.C. Leary, P.A. Cobine, F.B. Young, G.N. George, E.A. Shoubridge, and D.R. Winge, Human Sco1 and Sco2 function as copper-binding proteins. J Biol Chem 280 (2005) 34113–22.

[100] P. Buchwald, G. Krummeck, and G. Rodel, Immunological identification of yeast SCO1 protein as a component of the inner mitochondrial membrane. Molecular & general genetics: MGG 229 (1991) 413–20.

